# Proper control of R-loop homeostasis is required for maintenance of gene expression and neuronal function during aging

**DOI:** 10.1101/2021.06.29.450380

**Authors:** Juan Jauregui-Lozano, Spencer Escobedo, Alyssa Easton, Nadia A. Lanman, Vikki M. Weake, Hana Hall

## Abstract

Age-related loss of cellular function and increased cell death are characteristic hallmarks of aging. While defects in gene expression and RNA metabolism have been linked with age-associated human neuropathies, it is not clear how the changes that occur in aging neurons contribute to loss of gene expression homeostasis. R-loops are RNA-DNA hybrids that typically form co-transcriptionally via annealing of the nascent RNA to the template DNA strand, displacing the non-template DNA strand. Dysregulation of R-loop homeostasis has been associated with both transcriptional impairment and genome instability. Importantly, a growing body of evidence links R-loop accumulation with cellular dysfunction, increased cell death and chronic disease onset. Here, we characterized the R-loop landscape in aging *Drosophila melanogaster* photoreceptor neurons and showed that bulk R-loop levels increased with age. Further, genome-wide mapping of R-loops revealed that transcribed genes accumulated R-loops over gene bodies during aging, which correlated with decreased expression of long and highly expressed genes. Importantly, while photoreceptor-specific down-regulation of Top3β, a DNA/RNA topoisomerase associated with R-loop resolution, lead to decreased visual function, overexpression of Top3β or nuclear-localized RNase H1, which resolves R-loops, enhanced positive light response during aging. Together, our studies highlight the functional link between dysregulation of R-loop homeostasis, gene expression and visual function during aging.

## 1 INTRODUCTION

Aging is a process characterized by a time-dependent decline in physiological homeostasis that eventually leads to a loss of organismal function and increased incidence of death. Characteristic functional changes include loss of gene and protein expression, mitochondrial dysfunction, cellular senescence, and stem cell exhaustion (López-Otín et al., 2013). Aging is also a major contributor to development of many chronic diseases including ocular disease (Bonnel et al., 2003). Age-related vision loss and maculopathy have been associated with decreased density of retinal cells, including photoreceptors. Specifically, the age-related decline in photoreceptors affects predominantly rods rather than cones (Jackson et al., 2002). In addition, emerging evidence links age-related neurological diseases, including retinal neuropathies, to defects in gene expression and RNA metabolism (Parapuram et al., 2010). Nonetheless, the molecular mechanisms that contribute to the age-associated susceptibility of the eye to disease development are poorly understood.

R-loops are three-stranded nucleic acid structures consisting of an RNA-DNA hybrid and a misplaced single strand of DNA (Aguilera & García-Muse, 2012). They typically form during transcription in organisms ranging from yeast to humans and play a significant role in normal cellular physiology, being required for the initiation of mitochondrial replication and class switch recombination. Notably, recent studies suggest that R-loops can dynamically regulate gene expression (Niehrs & Luke, 2020); for example, due to their enrichment at gene termini, R-loops can modulate gene expression by preventing DNA methylation or limiting transcription factor access to promoters and facilitating efficient transcription termination at 3’-ends (Boque-Sastre et al. 2015; Skourti-Stathaki et al. 2014). Moreover, R-loop formation correlates positively with active transcription, gene length, GC content and DNA topology (Mackay et al., 2020).

Importantly, recent studies have shown that resolution of topological stress during transcription mediated by topoisomerases is critical for proper neuronal function (McKinnon, 2016). Topoisomerase 3β (Top3β) is a member of the type IA superfamily of topoisomerases, which unwind negatively supercoiled DNA formed during transcription and replication, an activity that prevents R-loop formation (Mackay et al., 2020). Loss of Top3β is associated with neurological disorders (Mackay et al., 2020) and has been shown to reduce lifespan in mice (Kwan & Wang, 2001).

Although R-loops are normal biological structures, their persistent formation is a major source of spontaneous DNA damage that can lead to transcriptional dysregulation and genome instability (Aguilera & García-Muse, 2012); two early hallmarks of aging. Given that aging is a main risk factor for development of chronic diseases, including neurodegenerative disease, it is conceivable that R-loops could play a significant role in age-associated mis-regulation of cellular functions. However, our understanding on R-loop biology during aging, and particularly in neurons, is quite limited. Due to high levels of transcription and alternative splicing, retinal cells and particularly photoreceptor neurons may be highly sensitive to RNA metabolism dysregulation. *Drosophila* compound eye contains approximately 800 units called ommatidia with each consisting of 20 cells, including eight photoreceptor (PR) neurons. The six outer PRs (R1-R6) expressing Rhodopsin 1 (Rh1) are mainly responsible for black and white vision and motion, and are similar to human rods. The inner PRs (R7 and R8) express Rh3/4 and Rh5/6, respectively, are responsible for color vision and resemble human cones. To characterize how aging impacts the genomic R-loop landscape in *Drosophila* photoreceptors, we isolated genetically labeled outer PRs using our recently improved nuclei immuno-enrichment method (Jauregui-Lozano et al., 2021) and performed MapR coupled with next generation sequencing (Yan & Sarma, 2020). Here, we show that R-loop levels in photoreceptor neurons increased progressively with age and were associated with genic characteristics, such as transcript levels and GC content. Further, our data show that majority of genes that decrease expression during aging contained R-loops. Finally, photoreceptor-specific depletion of DNA/RNA topoisomerase Top3β resulted in increased R-loop levels, reduced expression of long genes with neuronal function and reduced visual response. Importantly, overexpression of either Top3β or human RNASEH1, an enzyme that resolves R-loops, in the eyes resulted in enhanced visual response during aging. Together, our data show that aging is associated with increased levels of R-loops over transcribed genes, potentially disrupting transcriptional outcomes that contribute to age-associated changes in neuronal function, including visual response to light.

## 2 RESULTS

### 2.1 Aging photoreceptor neurons show increased global R-loop levels that correlate with loss of function and precede age-associated retinal degeneration

To examine the global R-loop levels in photoreceptor neurons, we tagged the outer nuclear membrane of R1-R6 with GFP fused to the KASH domain of Msp300 protein using the Rh1-Gal4 driver and isolated outer PR nuclei from the head homogenate with our tissue-specific method (Jauregui-Lozano et al., 2021). Next, we aged flies over the course of 50 days post-eclosion (emergence from the pupae) and extracted genomic DNA from isolated PR nuclei at three time points (Figure 1A). We then assessed the global R-loop signal with a DNA slot blot assay using the S9.6 antibody, which recognizes RNA-DNA hybrids. Specificity of S9.6 antibody towards RNA-DNA hybrids was shown by pre-treatment of DNA with ribonuclease H1 (RNase H1) that resulted in significant decrease of S9.6 signal (Figure 1B). Signal quantification showed approximately 30% increase in R-loop levels in PRs isolated from middle-aged, 30-day old flies as compared to that in young, 10-day old flies. This trend continued with a significant increase of nearly 50% in global R-loop levels at day 50 (Figure 1B-C). These data show that R-loops start accumulating early during photoreceptor aging, at a time point where flies show decreased visual function (Hall et al., 2017). Importantly, using optical neutralization, which measures photoreceptor structural integrity with light microscopy, we observed no retinal degeneration by middle age, with a stochastic loss of rhabdomeres occurring after day 40 (Figure S1). Thus, our data show that process of R-loop accumulation precedes age-related retinal degeneration, thus suggesting that increased R-loop formation may contribute to loss of function and possibly neuronal cell loss during aging.

**Figure 1.**
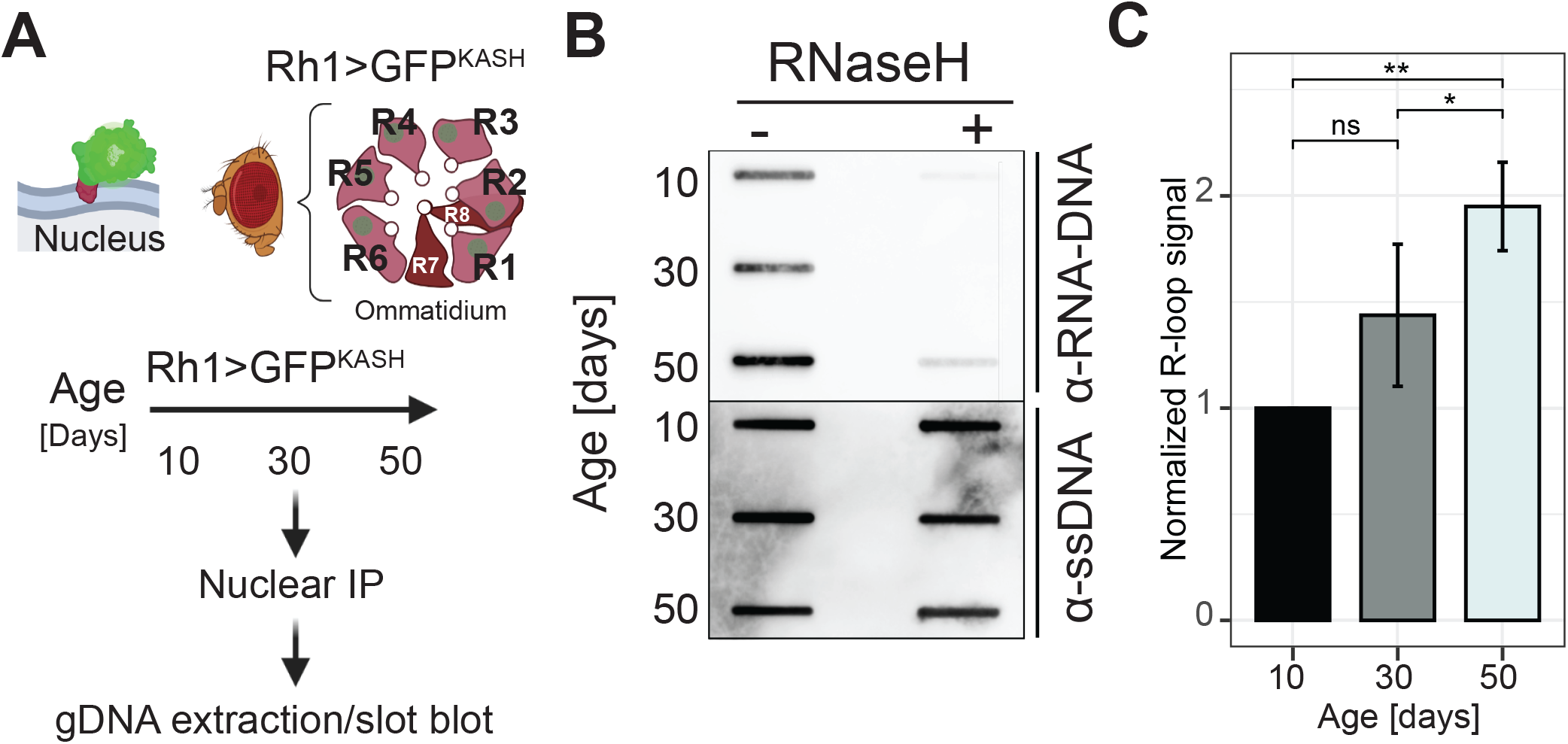
Aging photoreceptor neurons show increased global R-loop levels that correlate with loss of function and precede age-associated retinal degeneration. (a) Schematic of experimental outline to detect global levels of R-loops in aging photoreceptor neurons. Top: diagram of the cellular localization of the GFP^KASH^ protein. Dark blue lines represent each lipid layer within the nuclear membrane. Bottom: diagram of an ommatidium, a structural subunit in the *Drosophila* compound eye. Each ommatidium is composed of 8 photoreceptor neurons, labeled R1 to R8. Outer photoreceptors (R1-R6) express the *ninaE* (Rh1) gene. (b) Slot blot analysis of R-loop levels from photoreceptor nuclei at day 10, 30 and 50 post-eclosion treated with (right) or without (left) RNase H1. Slot blots were performed using RNA-DNA hybrid-specific S9.6 antibody (top) and ssDNA for loading control (bottom). (c) Quantification of S9.6 slot blot signal in aging PRs from (b). Values above 1 represent increase signal relative to day 10. Mean +/- Standard Deviation (SD), (n=2).

### 2.2 Profiling genome-wide R-loop distribution in PR neurons reveals age-associated changes

To determine the genomic R-loop landscape in aging PRs, we coupled our NIE approach with MapR, a recently published R-loop mapping strategy based on the specificity of RNase H1 enzyme to RNA-DNA hybrids, combined with the micrococcal nuclease (MNase)-based CUT&RUN technology. MapR uses a recombinant mutant form of MNase-fused RNase H1, which binds but does not degrade the RNA moiety within an RNA-DNA hybrid (ΔRH). Upon binding of the RNA-DNA hybrid by ΔRH-MNase, MNase activation by Ca^2+^ addition results in cleavage of surrounding DNA and a subsequent release of R-loop associated DNA fragment, which is used for library preparation coupled with high-throughput sequencing (Figure 2A). Surprisingly, we found that coupling NIE-purified photoreceptor nuclei with the standard MapR protocol yielded signal over genic regions resembling MNase-seq rather than R-loop specific enrichment. Our data showed MapR signal depletion around the Transcription Start Site (TSS) of genic regions (Figure S2A), suggesting that our samples were being over-digested by MNase. We therefore modified the standard MapR protocol based on the recently published improvement of the CUT&RUN method, which incorporates high salt with low calcium washing steps, and also decreased digestion time (see Methods in Supporting Information). To evaluate the quality of our modified protocol, we first compared our MapR data in *Drosophila* PRs to the original MapR data obtained in human HEK293T cells (Figure S2B) and found that the metagene profiles over the gene bodies showed similar R-loop distribution, with signal enrichment around the TSS and the Transcription Termination Site (TTS) (Figure S2A). In contrast, when we next compared our MapR data with R-loop mapping data obtained from *Drosophila* embryos using DRIP-seq (Alecki et al., 2020), we observed that DRIP-seq showed signal enrichment over gene bodies with a slight depletion around TSS (Figure S2C); discrepancies that have been shown previously to result from different affinities of S9.6 antibody used in DRIP-seq, as opposed to RNase H1 used in MapR, for RNA-DNA hybrids. In addition, the Alecki dataset was obtained during *Drosophila* early embryonic developmental stages. Thus, some differences in signal enrichment in our studies might arise from tissue-specific and method-specific effects.

**Figure 2.**
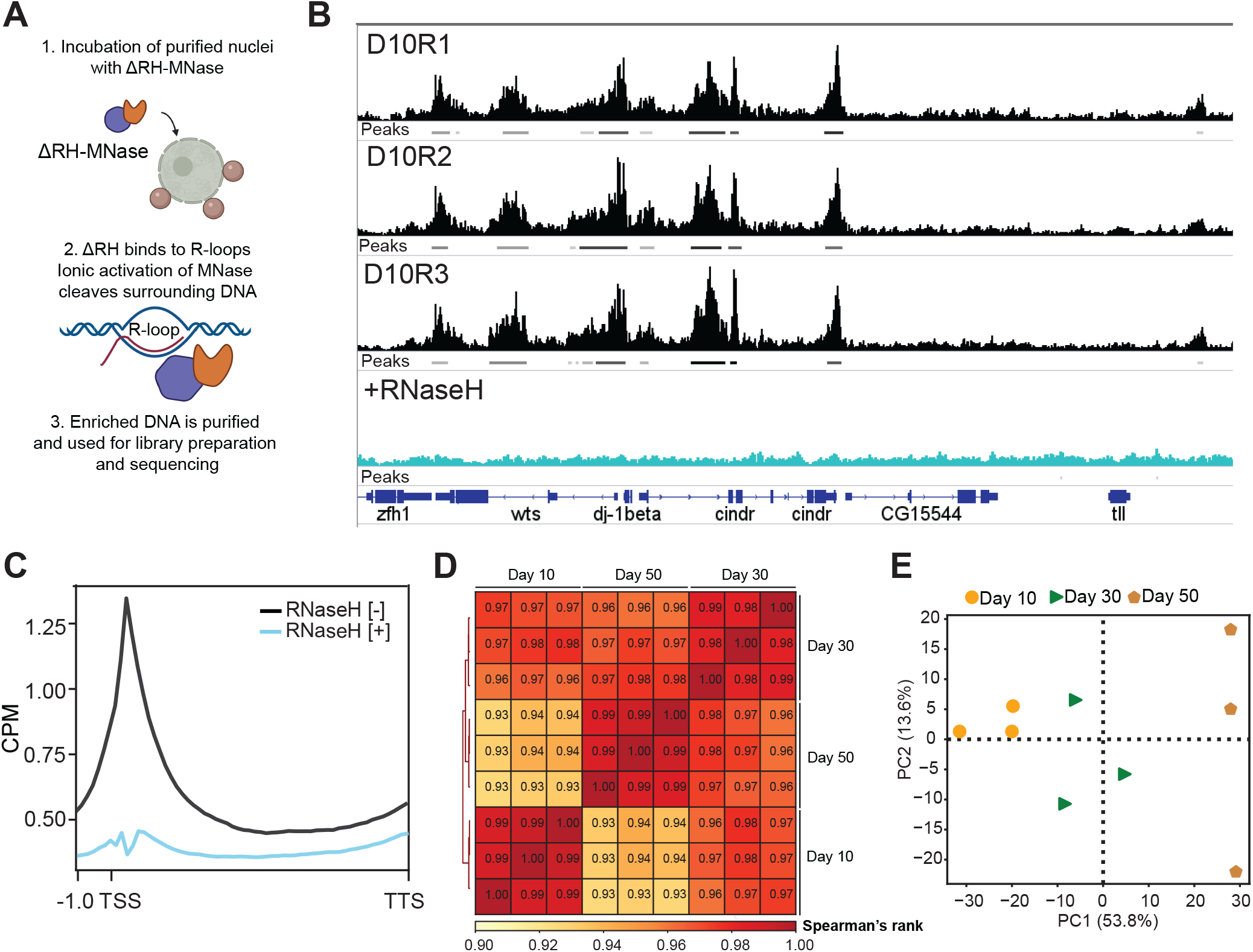
Profiling genome-wide R-loop distribution in PR neurons reveals age-associated changes. (a) Schematic diagram of the R-loop mapping technique used in this study (MapR). Immuno-enriched nuclei are incubated with ΔRNaseH1-MNase (ΔRH-MNase), where ΔRH binds to R-loops. Ionic activation of MNase results in cleavage of surrounding DNA and subsequent R-loop enriched DNA release, which is purified and used for sequencing library preparation. (b) Genome browser inspection of MapR track data on integrated genomic viewer (IgV) for a selected genomic region. Three independent biological replicates (R1-R3) from 10-day old flies’ samples not pre-treated with RNaseH1 are shown in black, and a sample from nuclei that were pre-treated with RNaseH1 prior to MapR (see Methods) is shown in blue. Peaks obtained using MACS2 for each sample are also shown as bars under each corresponding sample track. (c) Metaplot of CPM-normalized MapR signal over gene bodies for samples that were pre-treated with (blue) or without (black) RNase H1 prior to MapR (from b). (d) Spearman correlation heatmap of Aging MapR read distribution over 1000-bp binned genome. Scores between 0 and 1 shown in each box correspond to Spearman’s rank score. (e) Principal component analysis (PCA) of Aging MapR samples based on read distribution over 1000-bp binned genome.

To validate the specificity of our MapR method, we treated NIE-purified PR nuclei with RNase-H1 prior to performing MapR, which led to a complete loss of signal enrichment as shown by individual gene examples (Figure 2B) and gene metaplots (Figure 2C). In addition, when we normalized MapR signal by obtaining a ratio from RNaseH1 non-treated relative to treated samples, we found that the MapR signal distribution did not show significant changes in enrichment over genes bodies (Figure S2D). We note that the standard MapR protocol includes a separate negative control where nuclei are incubated with MNase alone to account for MNase binding. However, in our hands, the MNase control yielded no purifiable DNA, as shown by TapeStation profiles (Figure S2E).

Based on these observations, we performed MapR in aging PRs at day 10, 30 and 50 post-eclosion using our modified approach in three independent biological replicates which generated at least 3.5×10^7^ uniquely mapped fragments per sample (Supplemental File 1). Spearman’s correlation analysis based on read distribution over a 1000-bp binned genome revealed a strong positive association amongst the three biological replicates (Spearman’s p ≥ 0.96). In contrast, when we compared the samples between different age time points, we observed lower positive association between day 10 and day 50 (Spearman’s p ≥ 0.93), suggesting that the R-loop landscape changes were age-dependent (Figure 2D). Similarly, Principal Component Analysis (PCA) of the normalized R-loop distribution revealed that 53.8% variance amongst the samples for all biological replicates was attributed to age (Figure 2E). Notably, while samples clustered by age, the similarity amongst the biological replicates decreased with age (PC2 13.6%), suggesting that aging is associated with increased heterogeneity in R-loop distribution.

To further assess the quality of our sequencing datasets, we performed peak calling using the Model-based Analysis for ChIP-Seq (MACS2) algorithm using default settings and measured quality control metrics. To account for the differences in the number of mapped fragments in each sample (Figure S2F), we called the peaks using bam files that were down-sampled to the same number of mapped fragments (3.5×10^6^). Evaluation of the Fraction of Reads in Peaks (FRiP) score, which measures the quality of signal enrichment as defined by modENCODE, showed consisted FRiP scores higher than 0.37 for all samples (Figure S2G). Furthermore, we found that peak distribution was stably maintained during aging, with approximately 60% of peaks being annotated to promoters (TSS ± 2kb) and approximately 25% of peaks annotated to introns (Figure S2H), which is consistent with previously reported genome-wide R-loop distribution.

Taken together, our data demonstrate that we successfully applied the MapR method to tissue-specific samples in *Drosophila*, by purifying photoreceptor nuclei from the whole organism, and produced high quality R-loop mapping data. Furthermore, application of MapR in aging PRs showed that the genome-wide R-loop distribution changes in an age-dependent manner.

### 2.3 Age-associated R-loop accumulation over gene bodies correlates with high GC content, gene length and transcriptional levels

Given that R-loops typically form co-transcriptionally (Belotserkovskii et al., 2018), we next examined global and locus-specific distribution of age-associated changes in R-loops across actively transcribed genes, defined as having more than seven transcripts per million (TPM).

First, we analyzed global R-loop signal over gene bodies for actively expressed genes and compared the average signal across the gene, as counts per million (CPM). As expected, gene metaplots revealed that R-loop signal was enriched mainly over TSS and towards the 3’ ends of genes across all age time points (Figure 3A), which is consistent with previously reported R-loop distribution (Ginno et al. 2012; Sanz et al. 2016a; Nadel et al. 2015; Chen et al. 2017). Importantly, there was a widespread increase in R-loop levels over gene bodies during aging, most notably at day 50. Genome browser inspection of the averaged signal tracks for each time point for two individual genes showed changes in R-loop signal, including an early decrease or late increase (Figure S3A). To further evaluate the molecular characteristics of R-loop coverage during aging, we asked whether the age-associated increase in R-loops could be a consequence of broadening of the peaks. Consistent with our gene metaplots, MapR signal around the peaks showed that overall, R-loop coverage increased with age (Figure 3B), suggesting that R-loops might either extend or form at higher rate with age. To further assess an age-associated increase in R-loop occupancy, we quantified signal coverage, as defined by the sum of peak width for each time point. Notably, R-loop peaks covered approximately 18.7 megabases (Mb) of the genome at day 50, compared to 18.1 Mb at day 10, showing a modest but significant increase in coverage during aging (t-test, p<0.022) (Figure 3C). Supporting this data, violin plots depicting the peak width for all peaks revealed a slight but consistent increase in peak width at day 30 and day 50 as compared to that at day 10 (Figure S3C). Taken together, these observations show that R-loop signal over the genome accumulates with age, corroborating our findings from bulk R-loop levels using slot blots (see Figure 1). We note that the *Drosophila* genome has a total size of 180 Mb, and our data in *Drosophila* PRs showed that R-loops covered approximately 10% of the genome, which is similar to the genomic R-loop coverage obtained from other organisms, including mammals (Aguilera & García-Muse, 2012).

**Figure 3.**
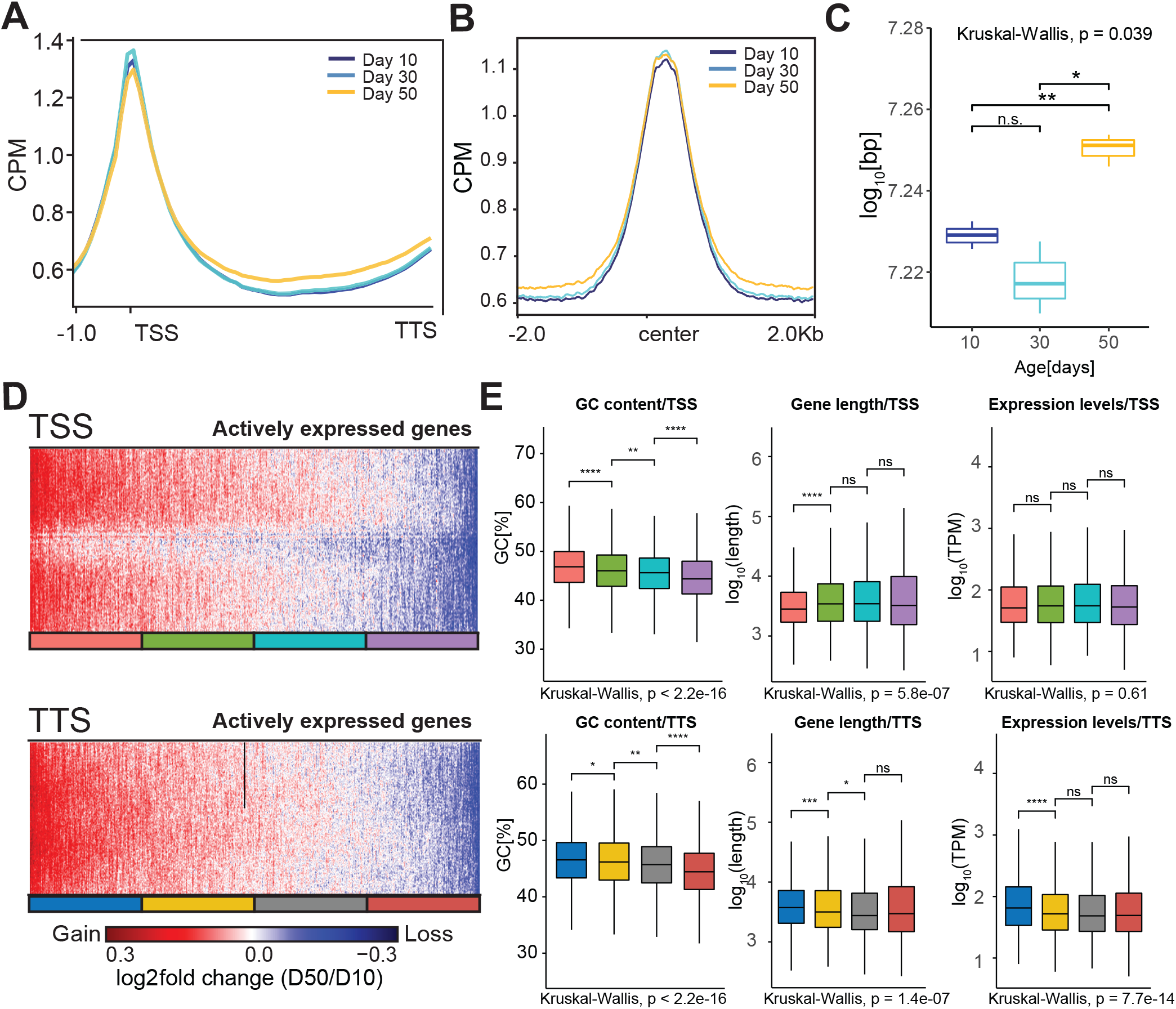
Age-associated accumulation of R-loops over gene bodies correlates with high GC content, gene length and transcript levels. (a) Metaplot of CPM-normalized MapR signal over gene bodies for all genes across age-timepoints. Signal is an average obtained from three independent biological replicates per age-timepoint. TSS indicates Transcription Start Site and TTS indicates Transcription Termination Site. (b) Metaplot of CPM-normalized Aging MapR signal around peaks obtained using MACS2 during aging. (c) Boxplot of genomic coverage of Aging MapR signal as defined by the total sum of peak width obtained at each time point. Peaks that mapped to scaffold or non-defined chromosomes were excluded from analysis. We used Wilcoxon Rank-Sum test to compare pair-wise differences in the distribution of genomic coverage amongst ages, (n=3). (d) Heatmap showing log2 ratios of Aging MapR signal around the TSS (top) or TTS (bottom), comparing day 50 to day 10. Genes are ranked based on their fold change value and divided in four groups (quartiles) based on their position on the heatmap. (e) Boxplot analysis of GC content, gene length and expression levels for each group of genes divided in four groups based on the Aging MapR fold changes around TSS (top) and TTS (bottom). p-value is obtained using Wilcoxon test.

To gain further insight into age-associated changes in R-loops, we compared the fold change in R-loop signal around TSS (±3 kb) or TTS (±3 kb) in old and young PRs. Heatmap plots of actively expressed genes ranked based on their fold change showed that the majority of the TSS-associated R-loops increased with age, while only approximately 50% of genes had an increase in R-loop signal around TTS (Figure 3D). Because R-loop formation is typically associated with specific genic characteristics such as gene expression level, torsional stress, and GC content (Chedin & Benham, 2020), we wondered if such genomic features were associated with an age-associated accumulation of R-loops over actively expressed genes. To test this, we clustered the genes from each heatmap from Figure 3D into four quartiles (Q1-4, with Q1 having the highest R-loop gain and Q4 having the highest R-loops loss), and assessed GC content, gene length and expression levels, respectively. Genes with TSS-enriched R-loops showed higher GC content than genes with R-loop losses (Wilcoxon-test, p<2.2×10^−5^), with no statistically significant association with long or highly expressed genes (Figure 3E-top). However, genes with TTS-enriched R-loops had high GC content, and were highly expressed and long (Kruskal-Wallis, p<4.5×10^−2^, p<4.5×10^−4^, and p<2.1×10^−9^, respectively) (Figure 3E-bottomC). Thus, these data show that age-associated accumulation of R-loops correlates with high GC content, length and expression levels.

### 2.4 Accumulation of R-loops in long genes correlates with decreased transcript levels in aging PRs

We previously showed that genes which decrease expression in aging PRs tend to be highly expressed and longer than genes that either increase or do not change expression (Hall et al., 2017). Moreover, aging *Drosophila* exhibit decrease in positive light response which correlates with decreased expression of long genes with neuronal function. Interestingly, recent reports have shown similar correlations between gene length and function in a variety of aging tissues and organisms, including humans (Lopes et al., 2021; Stoeger et al., 2019). Because our current data showed that long genes accumulated R-loops with age, and R-loops can lead to RNA polymerase II arrest and transcription inhibition (Bentin et al., 2005; Tous & Aguilera, 2007), we next investigated if there was an association between accumulation of R-loops and decreased gene expression during aging. Transcriptome profiling of PRs isolated from flies at day 10 and 50 revealed that 1700 genes (18%) were differentially expressed (DEG), with 722 genes (7%) decreasing expression and 978 genes (8%) increasing expression (p-adj<0.05, |FC|>1.5) (Figure 4A). To further evaluate the relationship between genic R-loop accumulation and gene expression, we identified R-loop containing genes (RCGs), by annotating high confidence peaks to the nearest TSS, and compared them to DEGs during aging. Venn diagram analysis revealed that 1388, or 69% genes were age-regulated at a transcript level and also contained at least one R-loop (RCGs/DEGs) (Figure 4B). Further, gene length analysis of each category showed that RCG/DEGs were significantly longer than either DEGs or RCGs alone (Figure 4C), suggesting that accumulation of R-loops in long genes could contribute to gene expression changes during aging (Wilcoxon test, adjusted p-value < 2.22e-16).

**Figure 4.**
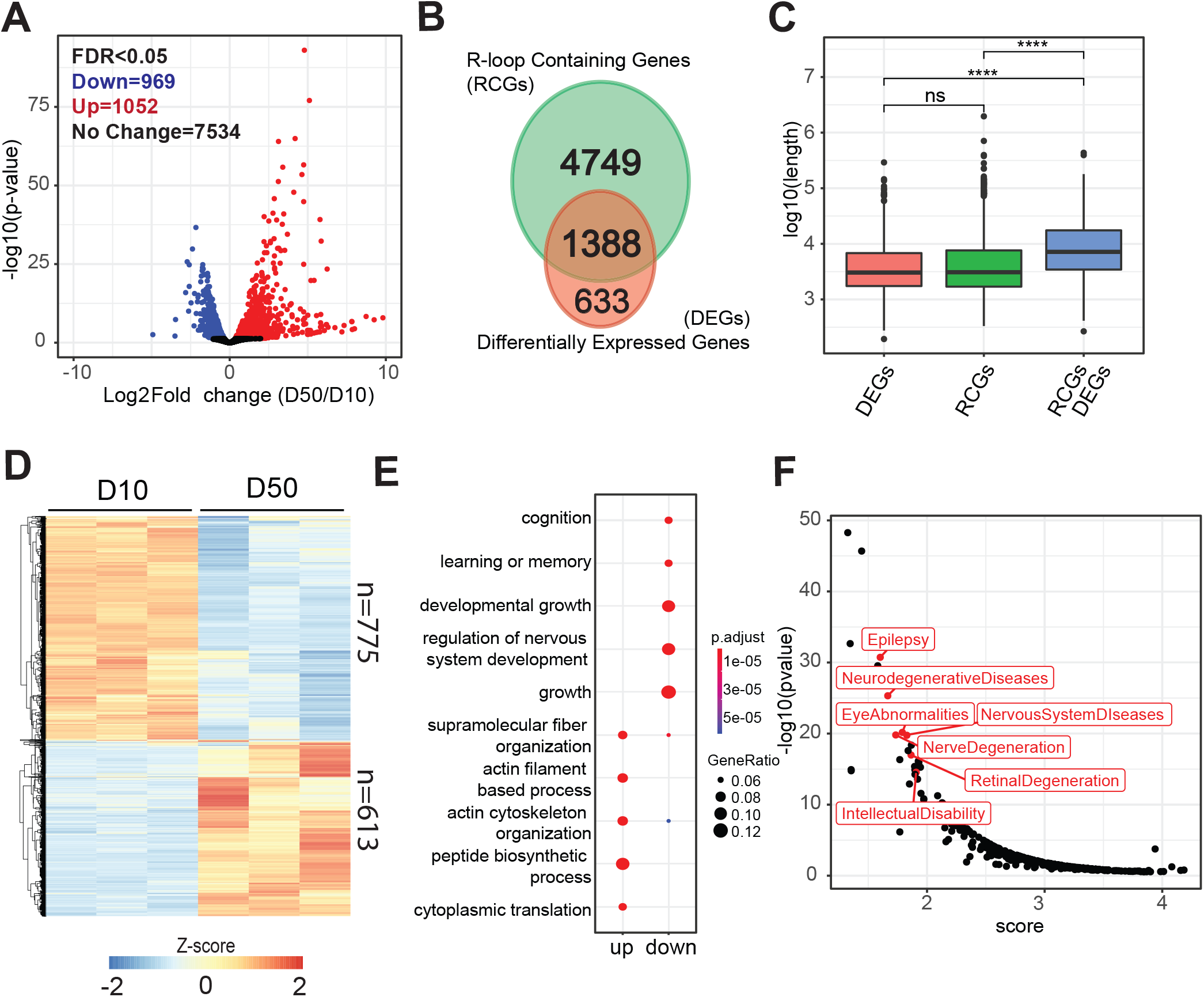
Accumulation of R-loops in long genes correlates with decreased transcript levels in aging PRs. (a) Volcano plot representing differentially expressed genes (DEGs) between day 50 and day 10. DEGs are obtained using DESeq2 (adjusted p-value < 0.05, |FC|>1.5). (b) Venn diagram representing the overlap between R-loop containing genes (RCGs) and DEGs from (a). (c) Box plot analysis of gene length for differentially expressed genes, R-loop containing genes, or RCG/DEGs from (b). p-value is obtained using Wilcoxon test. (d) Hierarchically clustered heatmap of RNA-seq data for RCG/DEGs from (b). Normalized Z-scores are calculated based on normalized counts obtained using DESeq2, and the heatmap is divided into genes that were either up- or down-regulated with age. (e) Dot plot of biological processes identified as significantly enriched in Gene Ontology (GO) term analysis for genes that were either up- or down-regulated from (d). (f) Scatter plot depicting an enrichment analysis of diseases associated with genes that were down-regulated with age and contained at least one R-loop. Analysis performed using literature mining tool BioLitMine (Hu et al., 2020). A lower score (x-axis) represents higher enrichment.

To gain further insight into the biological processes associated with RCG/DEGs, we first used hierarchical clustering and identified 613 genes (46%) to be up-regulated and 775 genes (54%) being down-regulated. Importantly, we found that overall, nearly 80% of the genes that decreased expression during aging accumulated R-loops, while only 58% of the genes that increased expression during aging contained R-loops (Figure 4D). Gene Ontology (GO) term analysis revealed that down-regulated RCG/DEGs were enriched for functional and neuronal categories, including cognition and regulation of nervous system development, whereas up-regulated RCG/DEGs were enriched for metabolic processes, such as peptide biosynthesis and translation (Figure 4E). To further characterize the RCG/DEGs, we used literature mining tool BioLitMine and identified medical subject heading (MeSH) terms associated with RCG/DEGs that were significantly enriched for several eye- and brain-relevant diseases, such as epilepsy, eye abnormalities, and retinal and nerve degeneration (Figure 4F). Collectively, our data showed that R-loops accumulated at both age up- and down-regulated genes, suggesting that R-loops may contribute to gene expression regulation via multiple mechanisms. For example, transcription can be blocked by direct collision of RNAP by pre-formed R-loops from previous transcription rounds or alternatively, can be inhibited by an intrinsic R-loop formation in the wake of ongoing RNAP. However, increased R-loop levels in age up-regulated genes may be simply a result of increased expression of these genes. Importantly, given that R-loops accumulated at most of the age down-regulated genes, which are enriched for long genes with neuronal function, it suggests that R-loops may contribute to regulation of biological pathways relevant for eye-specific functions.

### 2.5 Top3β depletion in *Drosophila* PR neurons leads to increased R-loop levels and decreased visual function

The neuronal transcriptome is enriched for long and highly expressed genes, that undergo high level of torsional stress during transcription (King et al., 2013; Liu & Wang, 1987). To solve DNA and RNA topological problems, cells use conserved topoisomerase enzymes that play a critical role in a wide range of fundamental metabolic processes in the genome (Pommier et al., 2016; Wang, 2002). One of the enzymes is Top3β, a highly conserved, dual-activity topoisomerase in animals that can change the topology of both DNA and RNA (Xu et al., 2013) and unwind negatively supercoiled DNA that forms during transcription, an activity that prevents formation of R-loops (Chedin & Benham, 2020). Loss of Top3β is associated with increased R-loop levels in mammalian cells (Yang et al., 2014) and has been shown to reduce lifespan in mice (Kwan & Wang, 2001). In addition, mutations in Top3β are linked to neurological disorders, thus highlighting the critical role of Top3β in neuronal function (Joo et al., 2020). Interestingly, our recent proteomic study of the aging *Drosophila* eye (Hall et al., 2021) revealed a 20% decrease in Top3β protein levels (Figure 5A, Figure S5A), suggesting that the aging eye might be sensitive to the loss of Top3β activity. Thus, we hypothesized that decreased Top3β levels could contribute to changes in R-loop homeostasis and neuronal function in aging PRs. To test this, we first depleted *Top3β* with ubiquitous RNAi in larvae (*tubP-Gal4>UAS-RNAi*) and measured bulk R-loop levels. Using DNA slot blot and RNA-DNA-specific antibody S9.6, we detected a 10% increase in R-loop levels in Top3β-depleted samples as compared to a control expressing non-specific RNAi (Figure 5B-C). Pre-treatment of DNA samples with RNase H1 led to a complete loss of the signal (Figure 5B-right), thus showing the specificity of the signal for RNA-DNA hybrids. In addition, qPCR analysis of Top3β transcript levels showed approximately 80% reduction in Top3β-depleted samples as compared to a control, thus validating the efficiency of the knockdown (Figure S5B). Taken together, these data show that Top3β in *Drosophila* has a conserved role in maintenance of R-loop homeostasis.

**Figure 5.**
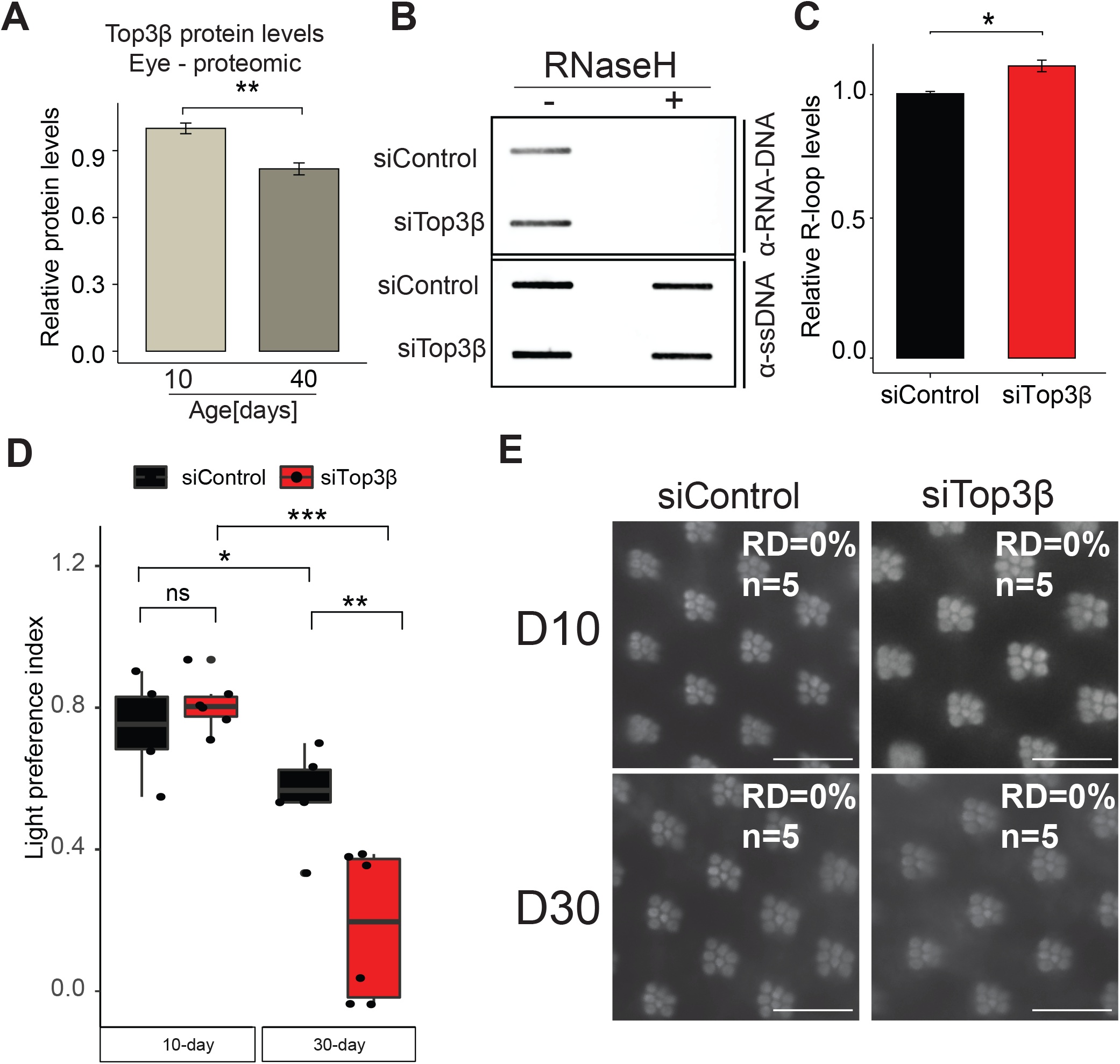
Loss of *Drosophila* Top3β leads to increased R-loop levels. (a) Comparison of Top3β protein levels in aging eyes from Rh1>GFP^KASH^ flies, shown as normalized protein abundance. Proteomic samples were prepared from 10- and 40-day old flies, 100 eyes/sample (n=4). Raw data taken from (Hall et al., 2021). (b) Slot blot analysis of R-loop levels from 3^rd^ instar larvae ubiquitously expressing siRNA against mCherry (siControl) or against Top3β (siTop3β). Samples were treated with (right) or without (left) RNase H1. Slot blots were performed using S9.6 antibody (top) and ssDNA for loading control (bottom). (c) Quantification of S9.6 slot blot from (b). S9.6 signal is normalized to ssDNA slot blot signal, (n=3). (d) Box plots showing the light preference indices (positive phototaxis) for Rh1>GFP^KASH^, mCherry-RNAi (siControl) or Rh1>GFP^KASH^, Top3β-RNAi (siTop3β) at day 10 and 30 (6 biological replicates for each time point or RNAi, 27 - 33 male flies/replicate; total number of flies per fly strain=150-180). p value obtained using Wilcoxon test. (e) Optic neutralization of siControl and siTop3β at day 10 and 30 post-eclosion from (d). Retinal degeneration (RD) scores were obtained by blindly quantifying 5 biological replicates. Score of 0% means there was no observable loss of rhabdomere or ommatidia.

Since loss of Top3β in *Drosophila* and mice leads to several neuronal phenotypes, such as disruption of synapse formation and behavioral impairments, we were next interested to see if depletion of Top3β specifically in PR neurons had any impact on visual function. Like most flying insects, *Drosophila* move towards light, thus exhibiting positive phototaxis (Choe & Clandinin, 2005). Importantly, we and others showed that positive phototaxis declines with age in flies (Carbone et al., 2016; Hall et al., 2017). To assess changes in visual behavior upon loss of Top3β, we depleted Top3β transcripts specifically in photoreceptors using *Rh1-Gal4>UAS-RNAi* and performed phototaxis assays at days 10 and 30 post-eclosion. As expected, there was approximately 15% decrease in positive phototaxis in the control flies between day 10 and day 30 (Figure 5D-Supplemental file 2). Notably, while flies with PR-specific depletion of Top3β showed no significant difference in the phototactic response at day 10, they showed approximately 60% decrease in visual behavior at day 30 as compared to a control (Wilcoxon test, p-value<0.47, and <0.015, respectively). Importantly, this decrease in visual behavior was not due to the loss of PR neurons, as optical neutralization showed no retinal degeneration in *Rh1-Gal4>UAS-RNAi* flies at day 30 post-eclosion (Figure 5E). Thus, our data show that Top3β is required for maintenance of proper visual function in aging *Drosophila* photoreceptor neurons.

### 2.6 Top3β regulates expression of a subset of long genes associated with neuronal function in photoreceptors

Given the role of Top3β in the resolution of torsional stress during transcription, we next hypothesized that Top3β might be required to regulate the expression of genes with neuronal function, which tend to be long and highly expresses (Zylka et al., 2015). To test this hypothesis, we analyzed the transcriptome of PR neurons depleted for Top3β in *Rh1-Gal4>UAS-RNAi; UAS-GFP*^*KASH*^ flies at day 30 using our NIE protocol. Differential expression analysis using DESeq2 between Top3β-RNAi and control, revealed that expression of approximately 1% of genes was regulated by Top3β (66 out 6500, FDR<0.05) (Figure 6A). Additionally, quantitative analysis of gene length based on whether a gene was differentially expressed in Top3β depleted PRs revealed, that genes with decreased expression were highly and significantly enriched for long genes relative to genes that either increased or did not change expression (Figure 6B). These data thus suggest that transcriptional regulation of long genes in the eye is particularly sensitive to decreased Top3β levels. Gene Ontology (GO) enrichment analysis of all Top3β-dependent genes revealed that genes with decreased expression were highly enriched for genes with neuronal functions as shown by the gene concept network analysis (Cnetplot) (Figure 6C). These genes included *Tenascin major* (*Ten-m; FBgn0004449*) and *Tenascin accessory* (*Ten-a; FBgn0267001*), which form a transmembrane heterodimer involved in synapsis regulation, *Tripartite motif containing 9* (*Trim9; FBgn0051721)*, a E3 ubiquitin ligase involved in neurogenesis, axon guidance, and eye development, and *knockout* (*ko; FBgn0020294)*, a storkhead-box protein involved in axon guidance. Importantly, GO term analysis of up-regulated genes did not lead to any significant biological category enrichment. Thus, our data show that in PR neurons, Top3β is required to maintain gene expression levels of long genes that are involved in neuronal function.

**Figure 6.**
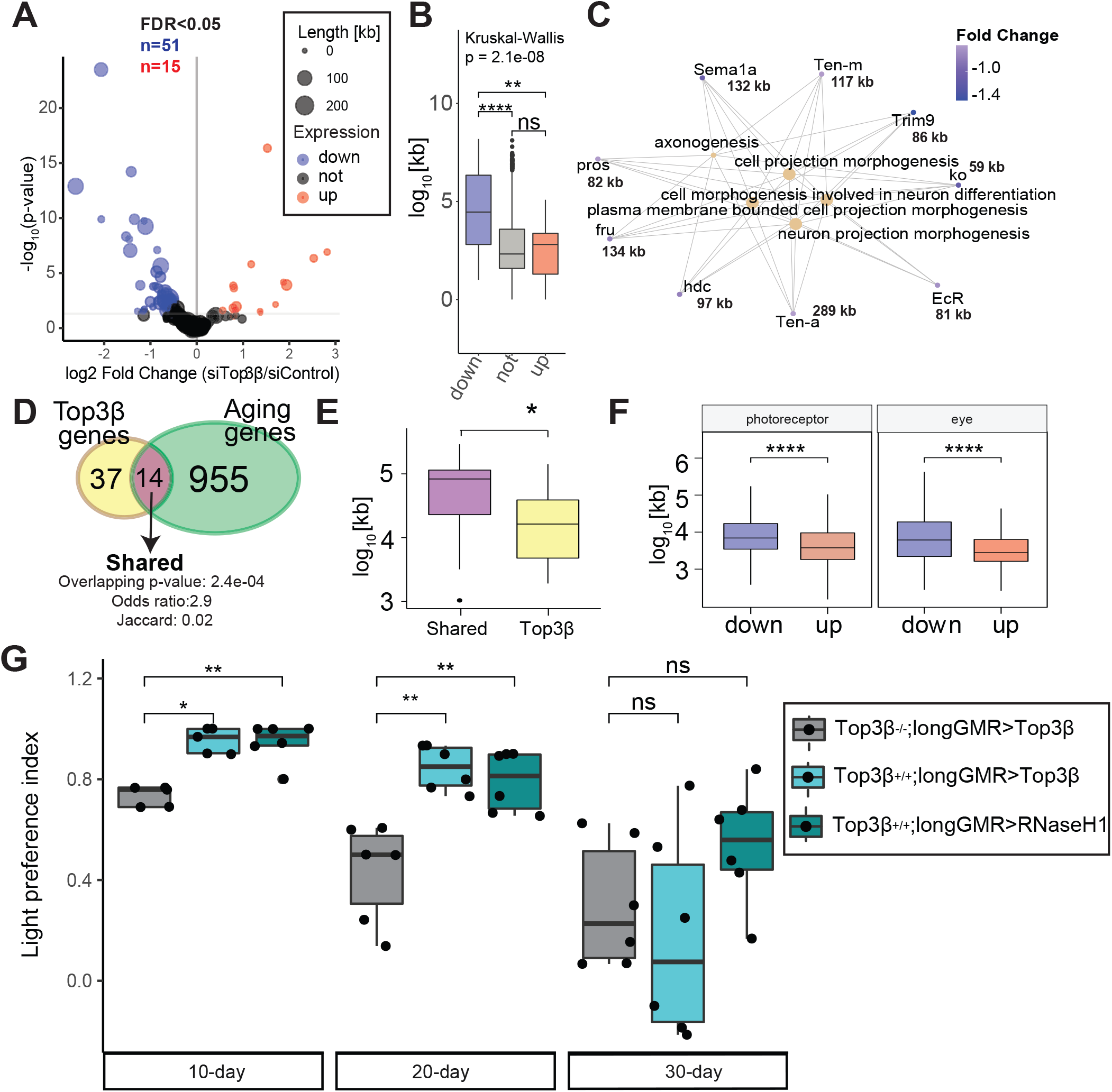
Top3β regulates expression of a subset of long genes associated with neuronal function in photoreceptors. (a) Volcano plot representing differentially expressed genes (DEGs) between siTop3β and siControl-expressing photoreceptors at day 30 post eclosion. DEGs obtained using DESeq2 (adjusted p-value < 0.05, |FC|>1.5). Size of each point reflect the gene length of the whole gene as defined as most upstream TSS and most downstream TTS. (b) Box plots showing the gene length (as log_2_-transformed bp) for genes identified as down-, up- or not regulated using DESeq2 in siTop3β photoreceptors relative to siControl (adjusted p-value < 0.05, |FC|>1.5). p-value obtained using Wilcoxon test. (c) Gene concept network analysis (Cnetplot) of genes downregulated in siTop3β photoreceptors relative to siControl. Gene length in kilobases is shown next to each gene. (d) Venn diagram representing the overlap of genes that were down-regulated in either aging (D50 vs D10) or upon loss of Top3β (siTop3β vs siControl). Overlap significance is denoted as a “overlapping p-value”, obtained with a hypergeometric test. Odds ratio and Jaccard index are measurements of similarity. (e) Box plots showing the gene length (as log_2_-transformed bp) for genes in the overlap identified in (d) or genes that were regulated by Top3β but not during aging. p-value is obtained using Wilcoxon test. (f) Box plots showing the gene length (as log_10_-transformed bp) for genes that were identified as down- or up-regulated in either aging PRs (left) or eyes (right). Eye data was obtained from (Stegeman et al., 2018). (g) Box plots showing the light preference indices (positive phototaxis) for Top3β-/-; longGMR>Top3β, Top3β-/-; longGMR>Top3β, and longGRM>RNaseH1 at day 10, 20, and 30 post-eclosion (6 biological replicates for sample; 29 - 31 male flies/experiment; total number of flies ∼180). p-value obtained using Wilcoxon test.

Next, we asked whether the expression of Top3β-dependent genes was mis-regulated during in aging PR neurons. To do this, we compared genes that were down-regulated either during aging or upon depletion of Top3β. Venn diagram revealed that 30% of Top3β-dependent genes were also down-regulated during aging (“shared genes”), with the overlap being statistically significant (Figure 6D). Further, individual inspection of these shared genes revealed that majority of them were long and showed similar changes in fold change expression during aging and Top3β depletion (Figure S6A). Thus, our data show that during aging, several long genes important for neuronal function decrease expression in a Top3β-dependent manner. Further, age- and Top3β-shared genes were significantly longer than Top3β-dependent genes (Figure 6E), suggesting that expression of longer genes is particularly sensitive to loss of Top3β during aging. However, gene length analysis of genes differentially expressed during aging in PRs revealed that down-regulated genes were significantly longer than up-regulated genes (Figure 6F-left). In addition, gene length analysis of our previously published RNA-seq data from aging eyes of Rh1>GFP^KASH^ flies (Stegeman et al., 2018) revealed a similar trend (Figure 6F-right) indicating that expression of the long genes in the aging retina is particularly sensitive to dysregulation of molecular mechanisms that include Top3β.

Collectively, our data showed that Top3β is required to maintain expression of a specific subset of genes with neuronal function that tend to be very long and thus are most likely sensitive to loss of topoisomerase activity due to high levels of torsional stress. This suggests that during aging, proper levels of Top3β are required to maintain R-loop homeostasis and expression of genes important for visual function.

### 2.7 Overexpression of Top3β or RNaseH1 enhances visual function during aging

To further validate the role of topoisomerase activity and maintenance of R-loop homeostasis in visual function during aging, we over-expressed either Top3β or human, nuclear localized, RNASEH1 in *Drosophila* eyes under the control of longGMR-GAL4 driver, which induces high expression in PRs and assessed visual behavior using phototaxis in 10-, 20-, and 30-day old flies. Importantly, we validated that RNASEH1 is expressed using qPCR and showed that bulk R-loop levels are decreased upon expression of RNASEH1 relative to a no driver control (Figure S6B-C-D). In addition, we over-expressed Top3β under control of longGMR in a Top3β null background (Top3β^-/-^;longGMR>Top3β). Our data showed that at day 10, over-expression of Top3β in the eyes of Top3β null flies resulted in similar light response as that of control flies (Rh1>siControl, see Figure 5D) or longGMR>siControl flies (Stegeman et al., 2018) (Figure 6G-Supplemental file 2). Furthermore, Top3β-/-; longGMR>Top3β flies showed an age-associated decrease in positive light response which is consistent with control Rh1>siControl flies (Figure 6G-grey). Importantly, flies with Top3β over-expression (Top3β^+/+^; longGMR>Top3β) showed an enhanced positive light response at either day 10 or day 20 (Figure 6G-light blue), corroborating that proper levels of Top3β contribute to visual response in flies. Moreover, over-expression of RNase H1 in *Drosophila* eyes (Top3β^+/+^; longGMR>RNaseH1) showed enhanced visual response in 10- and 20-day old flies as compared to control flies, similar to that of flies with Top3β over-expression (Figure 6G). These data suggest that Top3β function and R-loop resolution by RNase H1 contribute to proper visual function in *Drosophila*. Notably, qPCR analysis performed in flies with RNase H1 over-expression showed no significant difference in Top3β transcript level as compared to that in wild type flies (Figure S6E – purple vs blue) suggesting that the enhanced visual response in these flies is a result of increased R-loop removal. However, all tested genotypes showed decreased positive phototaxis at 30 days post-eclosion. Since optical microscopy imaging of the eyes did not show any changes in the eye morphology in 30-day old flies (Figure S6F), these data imply that additional aging mechanisms contribute to regulation of visual function in the eye, possibly including R-loops accumulated as a result of aberrant expression of additional R-loop metabolism associated factors.

## 3 DISCUSSION

While R-loops were previously considered to be mere byproducts of transcription, it has been demonstrated that R-loops play a significant physiological role in cellular biology of multiple organisms, including humans. Notably, there is a growing body of evidence that links R-loop accumulation to transcriptional imbalance and genomic instability, two main hallmarks of aging (López-Otín et al., 2013). Furthermore, dysregulation of R-loop homeostasis has been linked to human pathologies, including neurodegeneration (Groh & Gromak, 2014). Since age is the main risk factor for many neurodegenerative diseases, our current study focused on characterizing the changes in R-loop landscape induced during aging and evaluating the impact of R-loops on the gene expression, specifically in *Drosophila* photoreceptor neurons.

Characterization of global R-loop levels in aging PRs showed an increase in R-loops by middle age, with an additional significant increase late in aging. To our knowledge, this is the first published report demonstrating changes in R-loop levels in a specific tissue during aging. To further evaluate the R-loop distribution genome-wide, we modified a recently published R-loop mapping strategy, called MapR, coupled with high-throughput sequencing that can be useful when material is limiting. In addition, given that this method involves incubation of isolated nuclei/cells with a recombinant mutant form of RNase H1 tethered with MNase and therefore does not require modification of the organismal genome, it is well-suited for studies in whole animals. Using this approach, our data demonstrated that R-loops covered approximately 10% of the *Drosophila* PR genome, which is similar to the reported genomic distribution of R-loops in other organisms (Sanz et al., 2016; Wahba et al., 2016). Consistent with previous reports, R-loops were associated with known genomic hot-spots such as gene termini and specific genic features such as high GC content, gene length and expression level. The aging transcriptomes of multiple organisms and cell types show a positive correlation between transcriptional downregulation and specific genic features, such as gene length and GC content (Stoeger et al., 2019) and R-loops are known to play a key physiological role in transcription regulation due to their presence at promoters and terminators, where they regulate transcription initiation and termination, respectively (Niehrs & Luke, 2020). Thus, R-loop accumulation over these genomic regions may be a conserved mechanism that contributes to gene expression regulation in multiple cell types, including neurons.

Further, we observed a significant increase in R-loop signal over gene bodies and importantly, age-associated broadening of R-loop peak signal, suggesting that aging neurons accumulate R-loops at higher rate or R-loops are more persistent and potentially extend with age.

Transcriptome of neurons is biased for longer transcript relative to non-neuronal cell types (Zylka et al., 2015). Notably, long genes accumulate high topological stress during transcription and loss of topoisomerase activity has been shown to preferentially inhibit expression of (López-Otín et al. 2013) long genes (King et al., 2013). Since age-associated R-loop gains were particularly localized at long and highly expressed genes, we sought to further explore the impact of decreased topoisomerase activity on the maintenance of photoreceptor neuron homeostasis. DNA/RNA topoisomerase Top3β dysfunction is associated with increased R-loop levels in mammalian cells and mutations in Top3β are linked to neurological disorders, thus highlighting the critical role of Top3β in neuronal function (Joo et al., 2020). Importantly, Top3β protein levels decrease in aging *Drosophila* eyes (Hall et al., 2021). Here, we demonstrated that normal Top3β levels were required for maintenance of neuronal function, as shown by an age-associated decrease in visual behavior upon photoreceptor-specific downregulation of Top3β. In addition, depletion of Top3β in PRs lead to decreased expression of a subset of long genes with neuronal function, some of which also gained R-loop levels with age. Moreover, Top3β or RNase H1 over-expression in the eyes enhanced positive light response in flies and mitigated age-dependent loss of visual function. Collectively, our data suggest that Top3β may function in regulation of gene expression and maintenance of R-loop homeostasis in a subset of long genes required for neuronal function in aging photoreceptors. Thus, imbalance in R-loop homeostasis during aging could make postmitotic neurons particularly susceptible to dysregulation of gene expression and loss of function leading to increased risk of age-related neurodegeneration.

Aging is accompanied by elevated incidence of ocular diseases such as age-related macular degeneration and glaucoma, which exhibit characteristics of neurodegenerative diseases including loss of function and irreversible neuronal cell loss. How does aging impact development and progression of age-associated chronic diseases is one of the key questions in the biology of aging. Our current studies demonstrate a novel finding that *Drosophila* photoreceptor neurons progressively accumulate R-loops during aging, mostly at long and highly expressed genes. Importantly, integration of our transcriptomic and R-loop mapping data shows that majority of genes that decrease expression in aging PRs accumulate R-loops, thus suggesting that R-loops could be involved in cell physiology of aging neurons via inhibition of gene expression. Moreover, persistent formation of R-loops often leads to increased DNA damage, which is associated both with aging and neurodegeneration. Given that mutations in number of proteins involved in R-loop biology are implicated in neurodegenerative disease, our studies suggest that both aging and neurodegeneration may be sensitive to dysfunction in similar pathways.

## 4 EXPERIMENTAL PROCEDURES

See Supporting Information

## ACKNOWLEDGMENTS

Fly stocks from the Bloomington Drosophila Stock Center and information from FlyBase were used in this study.

## AUTHOR CONTRIBUTIONS

J.J-L. performed all the NGS experiments, bioinformatics analysis and figure generation. N.A.L. and V.M.W. assisted with data analysis. S. E. performed the phototaxis assay. A.N.E. and H.H. performed the slot blots and Western blots. J.J-L. and H.H. wrote the manuscript with input from other authors. H.H. supervised the project and conceived the study.

## DATA AVAILABILITY

RNA-seq expression data and MapR mapping data are accessible through Gene Expression Omnibus repository under series accession numbers GSE174488, GSE174491 and GSE174515, respectively. The following link has been created to allow reviewers to access these records while they remain in private status.

## HOW TO CITE THIS ARTICLE

Jauregui-Lozano J, Escobedo SE, Easton AN, Lanman NA, Weake VM, Hall H. Proper control of R-loop homeostasis is required for maintenance of gene expression and neuronal function during aging

## FIGURE LEGENDS

**Supplemental File 1**. Summary of uniquely mapped fragments for Aging MapR samples to *Drosophila melanogaster* dm6 genome.

**Supplemental File 2**. Raw data obtained in phototaxis assay for Figure 5D and Figure 6G

