## Supplementary figures and images for "Proper control of R-loop homeostasis is required for maintenance of gene expression and neuronal function during aging"

### Supplemental Figure S1

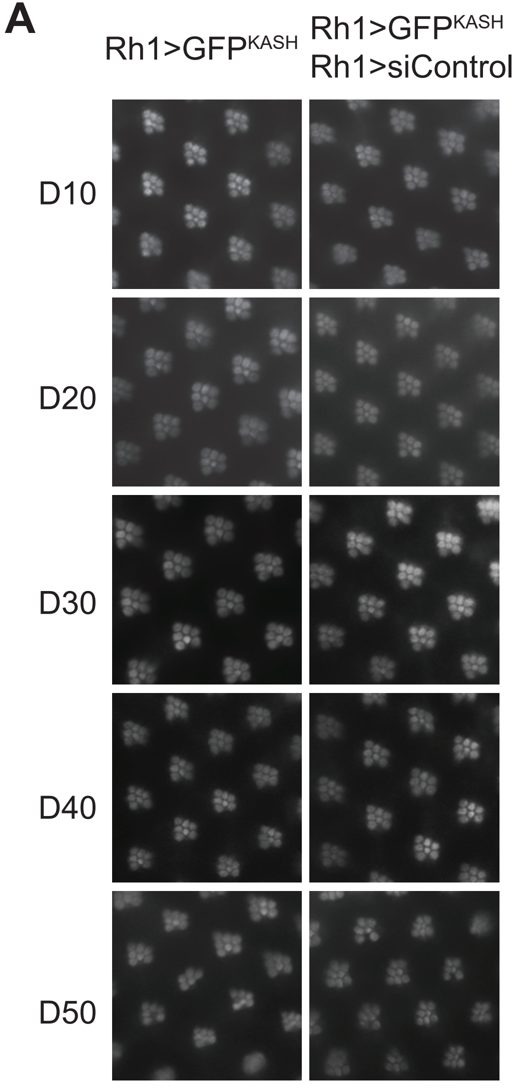

### Supplemental Figure S2

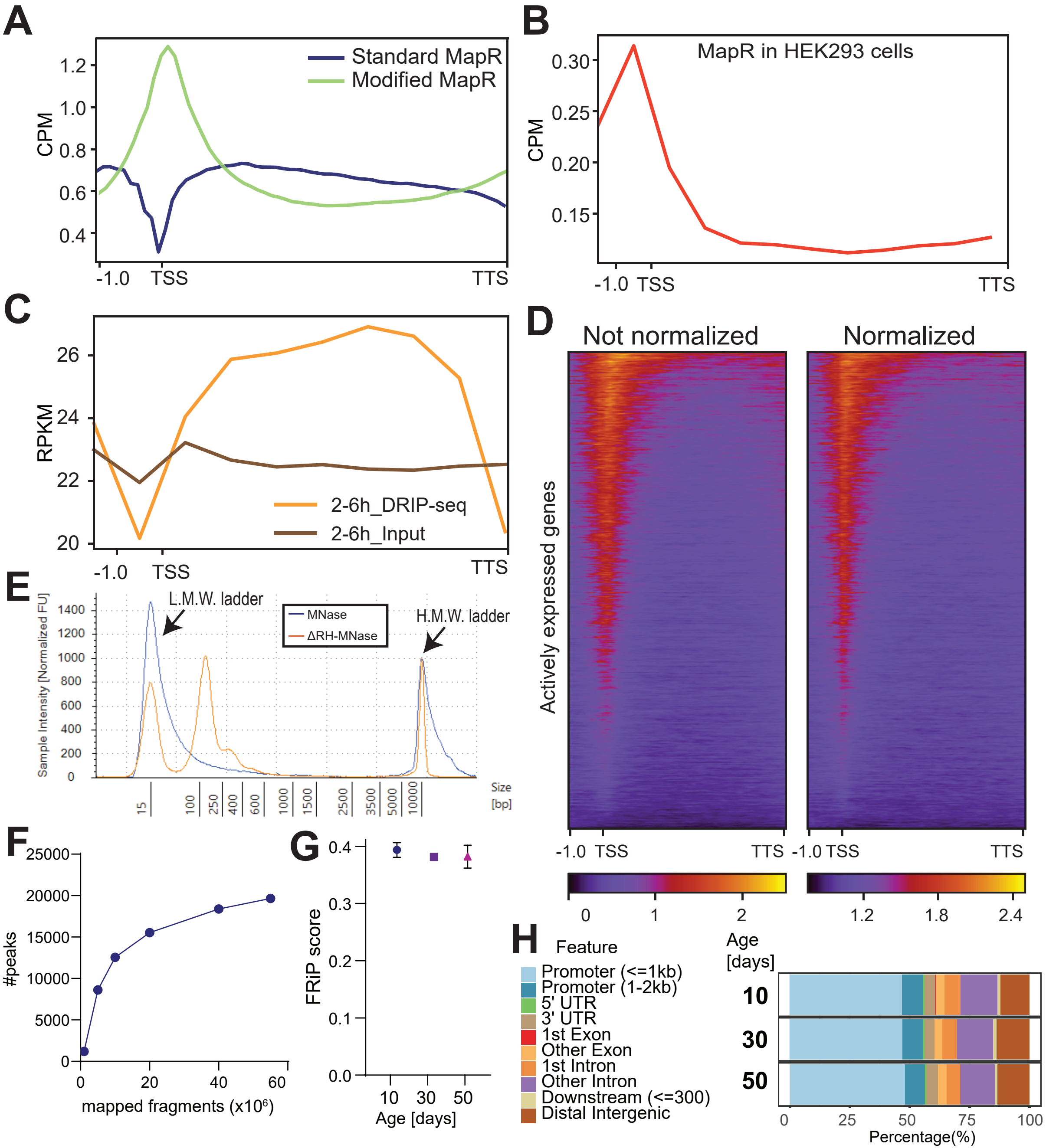

### Supplemental Figure S3

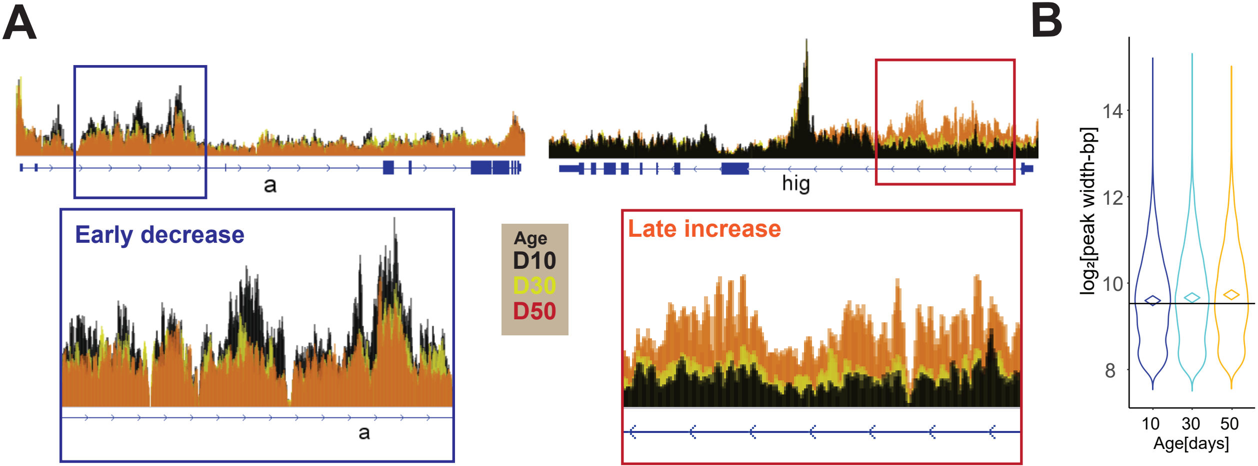

### Supplemental Figure S5

**A**

| M.S. (n=4) | | |
|---|---|---|
| Age | Abundance | Normalized |
| 10 | 120.6 | 1.01 |
| 10 | 113.3 | 0.95 |
| 10 | 126.5 | 1.06 |
| 10 | 116.5 | 0.98 |
| 40 | 105.7 | 0.89 |
| 40 | 99.3 | 0.83 |
| 40 | 91.2 | 0.76 |
| 40 | 94 | 0.79 |

**B**

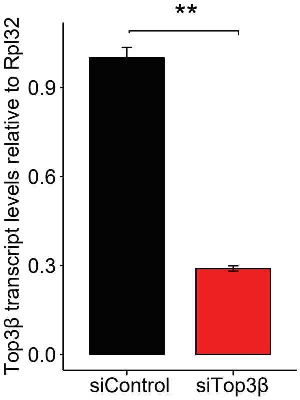

### Supplemental Figure S6

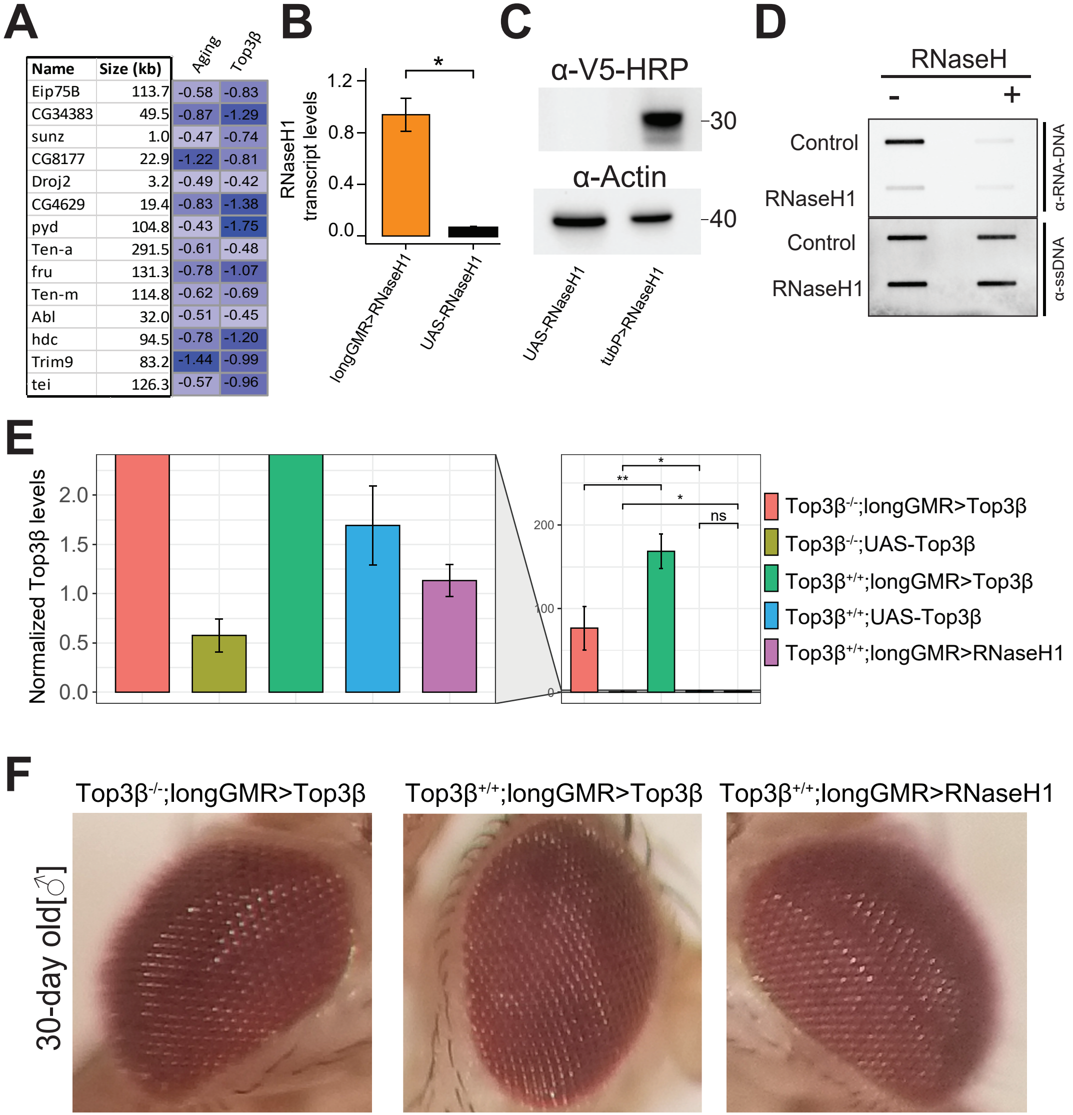
