## Supplemental Information for "Proper control of R-loop homeostasis is required for maintenance of gene expression and neuronal function during aging"

**Supporting information**

**4.1 *Drosophila* strains and fly maintenance**

Flies were raised in 12:12 h light:dark cycle at 25°C on standard fly food (Lewis, 1960). The following strains were used in this study: UAS-GFP^KASH^ (BDSC#92580), Rh1-Gal4 (BDSC#8691), longGMR-Gal4 (Bloomington Stock #8605), UAS-Top3β-RNAi (BDSC #31480), w1118; Top3β^26^;UAS-Top3βWT (were a gift from Weidong Wang), UAS-mCherry-RNAi (Stegeman et al., 2018). For siRNA experiments, UAS-Dcr2 was introduced in each genotype (BDSC#24645). To generate the UAS-RNaseH1-V5 flies, we used ppyCAG_RNaseH1_WT plasmid (Addgene #111906) to amplify the *Homo sapiens* RNASEH1 gene, including nuclear localization signal and the V5-tag, and cloned it as an EcoRI/NotI fragment into pUAST-attB plasmid (Bischof et al., 2007). Transgenic flies were generated using φC31-mediated transformation by BestGene (CA). For aging experiments, flies were collected over three days post-eclosion and maintained in population cages with a density of ~1000 flies/cage with fresh fly food change every two days. Male flies were aged to the specified age, collected and flash-frozen in liquid nitrogen at Zeitgeber time 6 (-/+ 1 hour).

**4.2 Nuclei Immuno-Enrichment (NIE) protocol**

NIE was performed as described previously (Jauregui-Lozano, 2021). Detailed protocol can be found at dx.doi.org/10.17504/protocols.io.buiqnudw. High-throughput experiments were done with three independent biological replicates, with each replicate consisting of 200-400 age matched male flies.

**4.3 Slot blot and Western blot**

DNA was extracted using Quick DNA MiniPrep Plus Kit (Zymo Research, Catalog #D4068) following the instructions for tissue homogenization. DNA concentration was measured using Qubit dsDNA settings. 35-50 ng of DNA was loaded on a Hybod-N^+^ membrane (GE Healthcare, Catalog #RPN119B) using slot blot apparatus. After drying, the membrane was blocked using 5% milk in PBST, then incubated with anti-S9.6 antibody in PBST (Millipore Sigma, Catalog # MABE1095), followed by goat anti-mouse HRP antibody (BioRad, Catalog *#170-6516*). For loading control, membranes were denatured in 0.5 M NaOH, 1.5 M NaCl for 10 min at RT, followed by 10 min-wash in PBST and neutralized in 1.5 M NaCl, 0.5 M TrisCl for 10 min at RT. The membrane was then washed with PBST for 10 min, blocked in 5% milk PBST for 30 min, and incubated with mouse anti single-stranded DNA antibody (MilliporeSigma, Catalog #MAB3034), followed by goat-anti-mouse HRP antibody (BioRad, Catalog *#170-6516*). To test the S9.6 antibody specificity, half of the DNA samples were treated with RNase H enzyme (NEB, Catalog #M0297S -10U) at 37^o^C for at least 3 HR or over/night. Western blotting analysis was performed using 50 µg of whole cell extract protein from larvae using the following antibodies: anti-V5-HRP (ThermoFisherScientific, Catalog #MA5-15367) and anti-actin (GeneTex; Catalog #GTX629630).

**4.4 GST-ΔRh-MNase and GST-MNase protein purification**

Protein overexpression and purification were performed as described previously (Yan & Sarma 2020). *Protein overexpression:* BL21 (DE3) competent *E. coli* cells (NEB, Ipswich, MA, Catalog #C2527H) were transformed with 10 ng of either pGEX-6p-1-GST-MNase or pGEX-6p-1-GST-ΔRNH-MNase plasmid (Addgene, Watertown, MA, Catalog #136291 and 136292, respectively). Transformed bacteria were grown in 500 mL of standard LB media (*LB (Luria-Bertani) Liquid Medium*, n.d.) supplemented with 100 μM Carbenicillin at 37^o^C. Once optical density reached 0.5 at 600 nm, protein expression was induced with 1 mM IPTG and cultures were grown at 37°C for additional 3 hours under constant rotation. Cells were pelleted using at 4ºC, 8000 RPM for 10 minutes. *Protein purification:* Bacterial pellets were resuspended in ice-cold 1X PBS buffer (ThermoFisher, Waltham, MA, Catalog #70011-044) and sonicated using a Branson Digital Sonifier in five 15sec-45sec ON/OFF cycles. Pierce™ Glutathione Magnetic Agarose Beads (Thermo Fisher, Catalog #78601) were used to purify GST-tagged MNase and RNAseH-MNase recombinant proteins according to manufacturer’s instructions.

**4.5 Modified MapR**

To profile genome-wide distribution of R-loops, we followed the MapR protocol (Yan et al., 2019) with some modifications according to the improved CUT&RUN protocol to decrease background MNase cleavage and digestion (Meers et al., 2019). Briefly, isolated nuclei were washed with 1 mL of Digitonin-containing wash buffer (20 mM HEPES-NaOH, 150 mM NaCl, 0.5 mM Spermidine, 0.02% Digitonin) freshly supplemented with EDTA-free cOmplete protease inhibitors (Sigma-Aldrich, St. Louis, MO, Catalog #11873580001). Nuclei were resuspended in 150 µl of Digitonin-containing wash buffer and 1 µM GST-ΔRh-MNAse or GST-MNase was added to a final concentration followed by one-hour incubation at 4°C with constant rotation. Nuclei were washed three times with 500 uL Digitonin-containing wash buffer, then washed one time with 1 mL of Low-Salt Rinse Buffer (20 mM HEPES, pH7.5, 0.5 mM spermidine, 0.05% Digitonin) freshly supplemented with EDTA-free complete protease inhibitor. Nuclei were resuspended in 200 µL of ice-cold calcium containing Incubation buffer (3.5 mM HEPES pH 7.5, 10 mM CaCl2, 0.05% Digitonin) and placed on wet ice for 60 seconds. Upon removal of supernatant, nuclei were resuspended in 150 µL of EGTA-STOP buffer (170 mM NaCl, 20 mM EGTA, 0.05% Digitonin, 20 µg/ml glycogen, 25 µg/ml RNase A), followed by 30-minute incubation at 37°C. DNA was extracted using Quick-DNA Microprep Kit (Zymo Research, Irvine, CA, Catalog #D4074). DNA was quantified with Qubit 1X HS DNA (ThermoScientific, Catalog#Q33203) and 2 ng of purified DNA was used to make sequencing libraries with Tecan Ovation Ultralow V2 DNA-Seq Library Preparation Kit-Unique Dual Indexes (Tecan, Switzerland, Catalog #9149-A01). Up to 16 libraries were pooled in one lane for paired-end 150 bp Illumina HiSeq sequencing.

**4.6 RNA-seq**

Isolated nuclei were resuspended in 100 μL TRI reagent (Zymo Research, Irvine CA, Catalog #R2050-1-200) and incubated at RT for 1 hour, followed by RNA extraction using the Direct-zol™ RNA Microprep (Zymo Research, Catalog #R2061). Purified RNA was quantified with the Qubit™ RNA HS Assay Kit (ThermoFisher, Catalog #Q32852) as per the manufacturers’ instructions**.** cDNA libraries were prepared with 10 ng of nuclear RNA using Ovation SoLo RNA-seq System including *Drosophila*-specific anyDeplete technology for rRNA depletion (Tecan, Redwood City, CA, Catalog #0502-32). Up to 16 libraries were pooled in one lane for paired-end 150 bp Illumina HiSeq sequencing.

**4.7 Quantitative PCR (qRT-PCR)**

cDNA was synthesized using 150-300 ng of RNA using EpiScript RNase H- Reverse Transcriptase (Lucigen, Middleton, WI, Catalog #ERT12910K). qRT-PCR was performed using Bullseye EvaGreen qPCR 2X master mix-ROX (Midsci, Valley Park, MO, Catalog #BEQPCR-R) or PR1MA qMAX Green qPCR mix No ROX (Midsci, Valley Park, MO, Catalog #PR2000-N-100) and using the following primers for *eIF-1a* forward 5'-GCTGGGCAACGGTCGTCTGGAGGC-3' and reverse 5'-CGTCTTCAGGTTCCTGGCCTCGTCCGG-3'; for *Top3β* forward 5'-GAATGGGCGCGCGGTCGGGTC-3' and reverse 5'-CGCATCAGTTCGACGGTGTTCAGTGCC-3', and for *RNaseH1 forward* 5’-AGGCATTAGACTTCCTGGGC-3’ and *reverse* 5’- TCCCTGCACTTGTCTTCCAC-3’.

**4.8 Optic Neutralization**

Flies were anesthetized on a CO_2_ pad and glued to a glass slide. Live rhabdomeres were imaged using brightfield light microscopy in an Olympus BX51 microscope, as described previously (Stegeman et al., 2018).

**4.9 Bioinformatic analysis**

Raw reads were trimmed using Trimmomatic version 0.39 (Bolger et al., 2014) to filter out low quality reads (Q>30) and clean adapter reads. Cleaned reads were aligned to the *Drosophila melanogaster* genome (BDGP Release 6 + ISO1 MT/dm6 from UCSC) using splicing-aware aligner STAR version 1.3 (Dobin et al., 2013) for RNA-seq and Bowtie2 version 2.3.5.1 (Langmead & Salzberg, 2012) for MapR using –sensitive settings. Samtools version 1.8 (Li et al., 2009) was used to obtain, sort and index BAM files. For genome browser inspection as well as further analyses, bigwig files were generated by normalizing datasets to count-per-million CPM coverage tracks with *deepTools* version 3.1.1 (Ramírez et al., 2014) using *--normalizeUsing CPM* settings. Spearman’s correlation scores were calculated using subpackages *multiBigwigSummary* and *plotCorrelation* as part of deepTools*.* Metaplots and genomic distribution heatmaps were generated with *computeMatrix, plotHeatmap* and *plotProfile* deepTool subpackages. MapR narrow peaks were obtained using MACS2 version 2.1.2 (Zhang et al., 2008) with standard settings. Peak overlap and genomic distribution of peaks was determined using R package ChIPseeker (v1.26.2) (Yu et al., 2015) and Homer (v4.11) Quantitative peak analysis was performed using GenomicRanges (v1.42.0) (Lawrence et al., 2013), csaw (v1.24.3) (Lun & Smyth, 2016) and edgeR (v3.13) (Robinson et al., 2010). R analysis was run in RStudio (v1.4.1106). Differential gene expression (DGE) analysis was performed using DESeq2 (Love et al., 2014). To increase the stringency of analysis, we generated shrunk fold change estimates within DESeq2 using the lfcShrink function (Zhu et al., 2019). DRIP-seq bigwig files from Drosophila embryos were downloaded from Gene Expression Omnibus GSE127329 (Alecki et al., 2020). Hierarchical clustering of gene expression and heatmaps were obtained using r package pheatmap (v1.0.12). Gene Ontology analysis was performed using clusterProfiler (v3.18.1). Gene overlap significance was calculated using GeneOverlap (v1.26.0). Mass Spectrometry counts for Top3β in the aging *Drosophila* eye were obtained from ProteomeXchange accession number PXD027090 (Hall et al., 2021)

**4.10 Phototaxis**

Phototaxis assay was performed as previously described (Stegeman et al., 2018). Briefly, 30 male flies were placed in a vial, tapped down and placed into the opening of the Maze apparatus. After letting them adjust to the darkness for 10 min, flies were given 30 seconds to choose between light and dark vial. Light preference index was then calculated as the proportion of flies that chose light minus the proportion of flies that chose dark. Flies were scored blindly.

**4.12 RNaseH treatment of NIE-purified nuclei**

Photoreceptor nuclei were purified using the NIE approach as described above, washed twice with 1X Dig-wash buffer, then resuspended in 45 uL of 1X Dig-wash buffer, and 5 uL of 10X RhBuffer. 4 uL (20U) of RNaseH (NEB, Catalog# M0297) enzyme was added and reaction was incubated for three hours at room temperature with constant rotation.

**4.13 Graph plots**

Bar-plots were generated using the R packages ggplot2 (v3.3.3) and ggpubr (v.0.4.0), and scripts used for RNA-seq analysis and plot generation are available upon request

**SUPPLEMENTAL FIGURE LEGENDS**

**Supplemental Figure 1**

(a) Aging optic neutralization time course of Rh1>GFP^KASH^ and Rh1>mCherry-RNAi, Rh1>GFP^KASH^, male flies at day 10, 20, 30, 40, and 50 post-eclosion. Retinal degeneration (RD) scores were obtained by blindly quantifying 5 biological replicates per genotype.

**Supplemental Figure 2**

(a) Metaplot of CPM-normalized MapR signal over gene bodies for all genes comparing the standard (Yan et al., 2019) and modified MapR protocol (current study) when protocol is performed using *Drosophila* photoreceptor nuclei.

(b) Metaplot of CPM-normalized MapR signal over gene bodies in HEK293T cells, originally from (Yan & Sarma, 2020).

(c) Metaplot of RPKM-normalized DRIP-seq signal in *Drosophila* 2-6h embryos, originally from (Alecki et al., 2020).

(d) Heatmaps of MapR signal from one biological replicate at day 10, when normalized using standard CPM (left) or normalizing to the CPM values obtained from RNaseH1-treated nuclei (right).

(e) Tape station profile from NIE-purified nuclei followed by treatment with ΔRH-MNase (orange trace) or MNase as negative control (blue trace). L/H.M.W. = Low/High Molecular Weight.

(f) Number of MapR peaks called using MACS2 based on sequencing depth. Bam file is down-sampled to 1, 5, 10, 20, 40 and 50 million mapped fragments and peaks are called.

(g) Fraction of Reads in Peaks (FRiP) scores for Aging MapR samples. Scores above 0.3 are commonly associated with high quality ChIP-seq datasets as defined my modENCODE standards.

(h) Genomic distribution of Aging MapR peaks. Promoter is defined as the region -2/+2 kb around TSS defined by RefSeq.

**Supplemental Figure 3.**

(a) Genomic browser inspection of two exemplar genes that showed an early decrease (left) or late increase (right) of R-loops during aging.

(b) Violin plots showing the R-loop peak width distribution for each age timepoint. Black line is placed at the calculated mean for peak width at day 10.

**Supplemental Figure 5.**

(a) Abundance and normalized abundance scores for Top3β-associated peptides (Hall et al., 2021) used to generate Figure 5A.

(b) Bar plots of quantitative PCR validating the downregulation of Top3β upon ubiquitous expression of an siRNA (BDSC#31480). p-value is obtained using t-test, (n=3).

**Supplemental Figure 6.**

(a) Table showing the genes identified in the overlap from (Figure6D), with the corresponding length and log_2_-transformed fold changes in gene expression identified in either aging (D50 vs D10) or upon loss of Top3β (siTop3β vs siControl).

(b) Bar plots of quantitative PCR analyzing transcript levels of RNaseH1 in longGMR>RNaseH1 (orange) or +;UAS-RNaseH1 (black) heads. p-value is obtained using t-test, (n=3).

(b) Western blot of TubP>RNaseH1 or UAS-RNAseH1 larvae.

(c) Slot blot analysis of R-loop levels from 3^rd^ instar TubP>RNaseH1 (referred as RnaseH1) or TubP>LacZ (referred as Control) larvae. Samples were treated with (right) or without (left) RNAse H1. Slot blots were performed using S9.6 antibody (top) and ssDNA for loading control (bottom).

(d) Bar plots of quantitative PCR analyzing transcript levels of Top3β in different samples as shown by the legend. p-values are obtained using t-test, (n=3).

(e) Representative images of eyes for genotypes tested in phototaxis (Figure 7) taken at 30-day old (n=3).
